# Molecular evidence of the eukaryotic parasites Perkinsea and Stylocephalidae in lizard muscle tissue

**DOI:** 10.64898/2026.10.17.759858

**Authors:** Erin S. Brown-Vera, Michael L. Yuan, Erin P. Westeen, Maggie R. Grundler, Ian J. Wang, Iris A. Holmes

**Author notes:** School Of Biological Sciences, Southern Illinois University, 1125 Lincoln Drive, Life Science II, Rm 389, Carbondale, Illinois 62901. Address correspondence to Iris A. Holmes.

## Abstract

Alveolates are a group of single-celled eukaryotes that include many important parasites of vertebrates, including the parasites that cause malaria. However, the group’s diversity and host associations in reptiles remain poorly understood. Here, we used 18S rRNA metabarcoding and phylogenetic analyses to identify novel parasites from tissue of iguanian lizards from the Caribbean and southern United States. This analysis revealed sequences closely related to members of the Alveolate families *Stylocephalidae* and *Perkinsea*, neither of which has previously been reported in reptiles. *Stylocephalidae* sequences clustered with gregarine parasites typically associated with arthropod hosts, while *Perkinsea* sequences grouped with environmental and freshwater lineages related to known amphibian pathogens. These findings represent the first molecular evidence of *Stylocephalidae*- and *Perkinsea*-related DNA in lizard tissues and suggest previously unrecognized ecological interactions between reptiles and alveolate microorganisms. Although infection cannot yet be confirmed, our results highlight the need for targeted environmental sampling, histopathology, and experimental studies to clarify host range, transmission potential, and possible conservation implications.

## INTRODUCTION

Over the past decades, reptilian diversity has declined, with 21% of reptile species currently threatened with extinction (Cox, 2022). A portion of this loss is attributable to diseases, some of which may be caused by unknown or emerging pathogens and parasites (Allain & Duffus, 2019). However, because reptilian parasites are less studied than those of other vertebrate groups (Tompkins et al., 2015), the diversity of known and emerging parasites in reptiles remains unclear. Reptiles fill important niches in many ecosystems as predators and prey, and provide important ecosystem services such as seed dispersal (Berriozabal-Islas et al., 2017), pollination, and pathogen regulation (Valencia-Aguilar et al., 2013). Therefore, understanding the diversity and potential conservation impact of parasites in reptile populations is crucial for maintaining species stability and whole ecosystem health (Tompkins et al. 2015).

Identifying new or emerging parasites before they cause conservation-relevant declines can be difficult. Microorganisms that are typically non-pathogenic or cause limited disease in coevolved host populations can nevertheless become major conservation threats through either the evolution of increased virulence or introduction into naïve host communities lacking resistance (Samuel et al., 2015; Peeler et al., 2010). The amphibian chytrid fungus *Batrachochytrium dendrobatidis* (Bd) exemplifies both processes: while endemic Bd lineages have long persisted in some host populations with limited pathogenic effects due to host–pathogen coevolution, the emergence and global spread of particularly virulent lineages have resulted in severe disease outbreaks and widespread population declines or extinctions in susceptible hosts (Belasen et al., 2022). This combination of evolutionary change and global pathogen spread highlights how previously harmless microorganisms can rapidly become devastating emerging pathogens. Given this possibility, it is important to survey unknown parasite diversity before it becomes a conservation issue.

Alveolates are among the most frequently reported reptile parasites (Jacobson, 2007), and previous genetic studies have shown significant diversity among alveolates in reptiles (Tomé et al., 2020). Alveolates are a taxonomic supergroup within Eukaryota that includes the Apicomplexan parasites. Although the Alveolata supergroup includes multiple serious parasites of vertebrates, including *Toxoplasma gondii*, *Cyclospora cayetanensis*, and *Babesia microti*, the full diversity of Alveolata is poorly understood. The group comprises over 6,000 described species and potentially thousands of unidentified species (Votýpka et al., 2017). One of the most well-known diseases caused by an alveolate is malaria (Okepeku et al., 2022), caused by members of the genus *Plasmodium.* Although it is most widely known as a human disease, different species of *Plasmodium* infect diverse hosts, including lizards such as *Anolis* (Telford et al., 1989) and *Sceloporus* (Motz et al., 2014). Other alveolate genera have also been reported in reptiles, including members of the genus *Hepatozoon* (Maia et al., 2014). These well-known parasites are known to have complex, often multi-host life cycles and transmission modes (Plattner & Soldati-Favre, 2008). For example, *Plasmodium* and *Hepatozoon* parasites require invertebrate hosts to complete their life cycle and are transmitted by arthropod vectors (Sato, 2021; Paperna et al., 2002).

We analyzed apparently novel 18S rRNA gene sequences obtained from muscle tissue samples and compared them to sequences from NCBI GenBank. Using phylogenetic approaches, we evaluated the relationships between our novel sequences and known alveolate lineages to infer their likely taxonomic placement and evolutionary history. Given the limited understanding of alveolate diversity in reptiles and the potential for pathogen emergence, this work will inform our understanding of the presence and diversity of understudied microorganisms in reptiles and highlight potential conservation concerns regarding these microorganisms.

## MATERIALS AND METHODS

### DNA Extraction and Amplification

Samples were collected from New Mexico, Florida, California, Alabama, Arizona, Sint Eustatius, or the collections of the Cornell Museum of Vertebrates and the University of Florida (Figure 1). The genera included in this study were *Sceloporus* (*S. cowlesi, S. woodi, S. occidentalis, S. virgatus*), *Anolis (A. carolensis, A. schwartzi*)*, and Aspidocelis* (*A. exsanguis*) (Supplemental Table 1). To extract alveolate DNA from squamate specimens, we extracted total DNA from tail and liver samples using Qiagen DNeasy Blood and Tissue extraction kits, following the manufacturer’s protocol. We then used endpoint PCR with primers designed by Wahab et al. (2020) and Chaudhry et al. (2019) to amplify the alveolate 18S rRNA gene, a conserved molecular marker widely used for the detection, identification, and phylogenetic placement of microbial eukaryotes (Pawlowski et al., 2012). The forward primer for all Alveolate taxa was 5’-GAGGTAGTGACAAGAAATAACAATA-3’. We used two reverse primers, one optimized for *Theileria* and *Babesia* and related taxa (including *Hepatozoon);* 5’-TCTTCGATCCCCTAACTTTC-3’, and one optimized for for *Plasmodium* and related taxa;5’-TATCTGATCGTCTTCACTCCCTTAAC-3’.After PCR amplification, we ran gel electrophoresis to detect the amplification of fragments between 400-500 base pairs long. Samples were Sanger sequenced with their respective forward primers at the Cornell University Biotechnology Resource Center. These sequences are available in GenBank under accession numbers PX926116-PX926129. Sequences were trimmed in SnapGene at the beginning and end of reads to remove ambiguous bases. We queried our sequences to the NCBI nucleotide database using the BLAST online tool. We discovered 14 sequences that did not match to the known alveolate parasites of reptiles, such as *Plasmodium* or *Hepatozoon*. We used these sequences for phylogenetic analyses.

**Figure 1:**
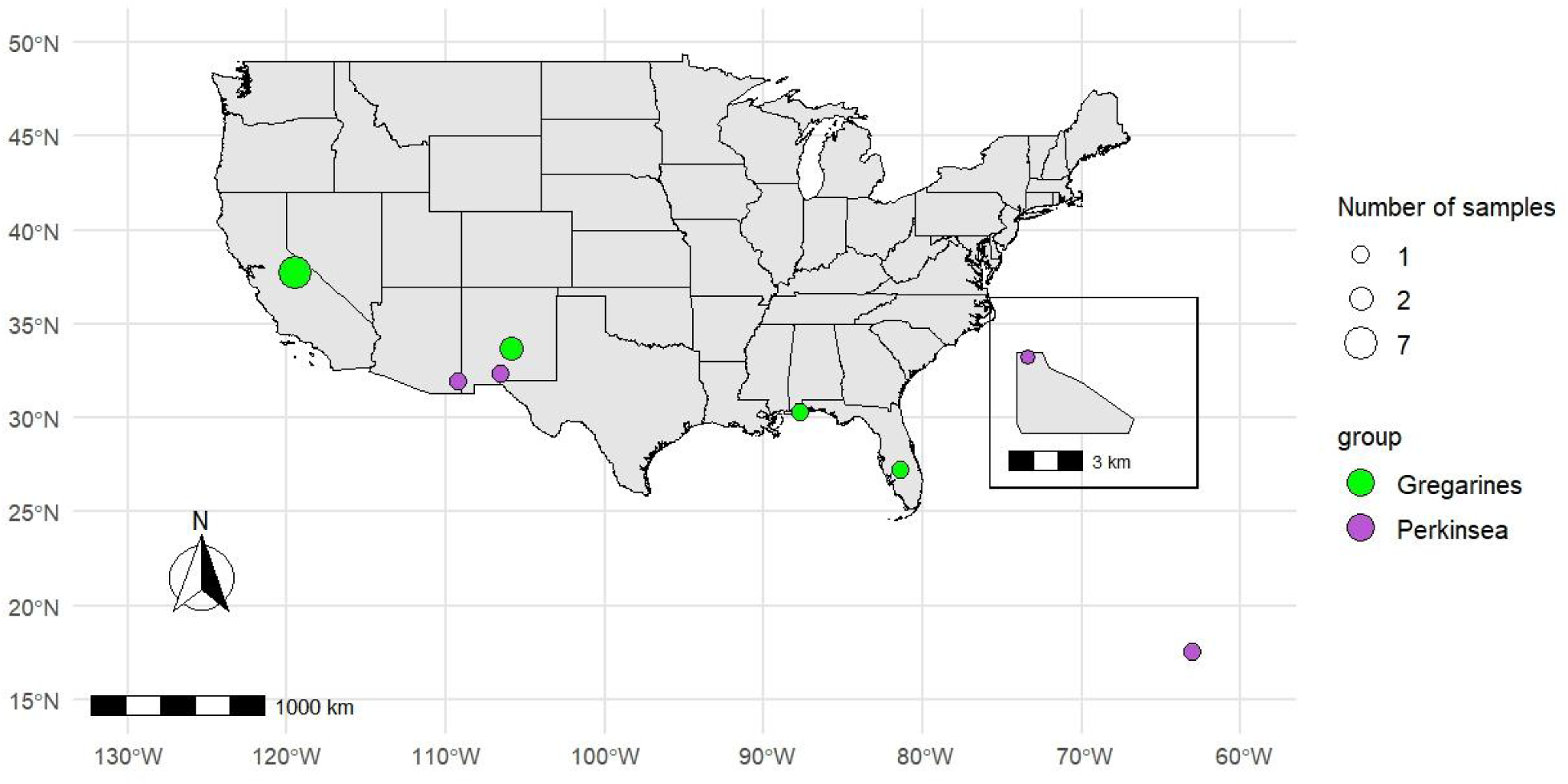
Map of Sample locations with novel parasites. Map showing the locations of samples with novel parasites across the mainland United States and the Caribbean island of Sint Eustatius (inset). Samples are colored by parasite group, as identified by BLAST comparisons. Scale bar and north arrow are included.

### Phylogenetic placement of unknown alveolate sequences

We downloaded the three top hits produced by the NCBI Blast algorithm, ranked by e-value. In all of our samples to include in our phylogenetic analyses, the highest e-value was 1.00E -37 (Supplemental Table 2). We supplemented these sequences with a previously published database of free-living and host-associated apicomplexan sequences, chosen to represent diverse evolutionary lineages (del Campo et al., 2023). Gregraine-related analyses included 50 reference 18S rRNA gene sequences (33 BLAST matches to our sequences and 17 additional diverse references) from 40 diverse members, including those known to infect reptiles (Supplemental Table 3). Perkinsea-related analyses included 88 reference 18S rRNA gene sequences (nine BLAST matches to our sequences and 79 additional diverse references), including amphibian-associated sequences (Supplemental Table 4).

All sequences were aligned using MUSCLE v5.1 (Edgar, 2021) through the **msa** R package (Bodenhofer et al., 2015). The alignment was exported in NEXUS format using **ape** (Paradis & Schliep, 2019). Bayesian phylogenetic inference using the GTR+Γ substitution model (nst=6, rates=gamma) was performed in MrBayes v3.2.7 (Ronquist et al., 2012). GTR+Γ was chosen because it is general, flexible, and accounts for both differing substitution rates between nucleotides and rate variation across sites, which is ideal for most nucleotide sequence datasets. Four MCMC chains were run for 1,000,000 generations, sampling every 500 generations, and discarding the first 25% of trees as burn-in. Convergence was verified by monitoring the average standard deviation of split frequencies (<0.01). The resulting consensus tree was imported into R using the **treeio** package and visualized with **ggtree** (Yu et al., 2017). Trees are presented in both pruned (with clades distantly related to our sequence removed for ease of visualization) and full, unpruned forms.

## RESULTS

The best matches to our unknown alveolate sequences from the NCBI database were within the taxonomic groups of Stylocephalidae, a family within the phylum Apicomplexa, and Perkinsea, a class within the monotypic phylum Perkinsozoa. Both taxa are in the supergroup Alveolata.

In order to identify the eleven parasite sequences matching gregarine taxa not previously reported in reptiles in BLAST comparisons, sequences from all gregarine groups in our reference dataset were used to make a phylogenetic tree. Phylogenetic analysis found parasite sequences amplified from our lizard samples to be most closely related to taxa typically isolated from insects, rather than parasites associated with squamates or other vertebrates (Fig. 2, Fig. S1). Sequences from our squamate samples were all nested within a clade comprising the family Stylocephalidae. All of these Stylocephalidae reference sequences came from beetle or silverfish hosts. The sequence from sample ARCH19 (*Sceloporis woodi* from Archbold Research Station in Florida) was placed sister to an *Atacamagregarina paposa* sequence isolated from a darkling beetle (*Scotobius brevipes*) with a posterior probability of 0.67 (Fig. 2). The sequences from samples E271 and EPW147 (both *Sceloporus cowlesi* from New Mexico) were identical and were placed with a posterior probability of 0.91 into a clade containing members of the genera *Stylocephalus* and *Xiphocephalus* (which in some studies (Mondal & Modak 2026, Nitsche et al. 2023) have been suggested to be the same genera). The reference sequences from these genera were derived from darkling beetle hosts (*Eleodes opaca, Eleodes obscura, Eleodes tricostata, Scaurus punctatus, Blaps parvicollis*, *Psectrascelis intricaticollis ovata*) (Fig. 2). The sequences from samples RCT674B (*Anolis sagrei* from California), and IW3137, IW3132, IW3136, IW3126, IW3135, IW3125, and IW3138 (*Sceloporus occidentalis* from California) formed a clade sister to a *Colepismatophila watsonae* sequence from a silverfish (*Lepisma saccharinum*) host, with a posterior probability of 0.97 (Fig. 2). All of the included reference sequences that came from other reptiles were not closely-related to our samples (Fig. 2). Together, this suggests that the gregarine sequences from our lizard samples are more closely related to insect parasites in the family Stylocephalidae than they are to previously described vertebrate parasites. We note that these novel reptile sequences came from lizards from across the southern United States, ranging from California to Florida, suggesting a potentially widespread association of Stylocephalidae gregarines and squamates.

**Figure 2:**
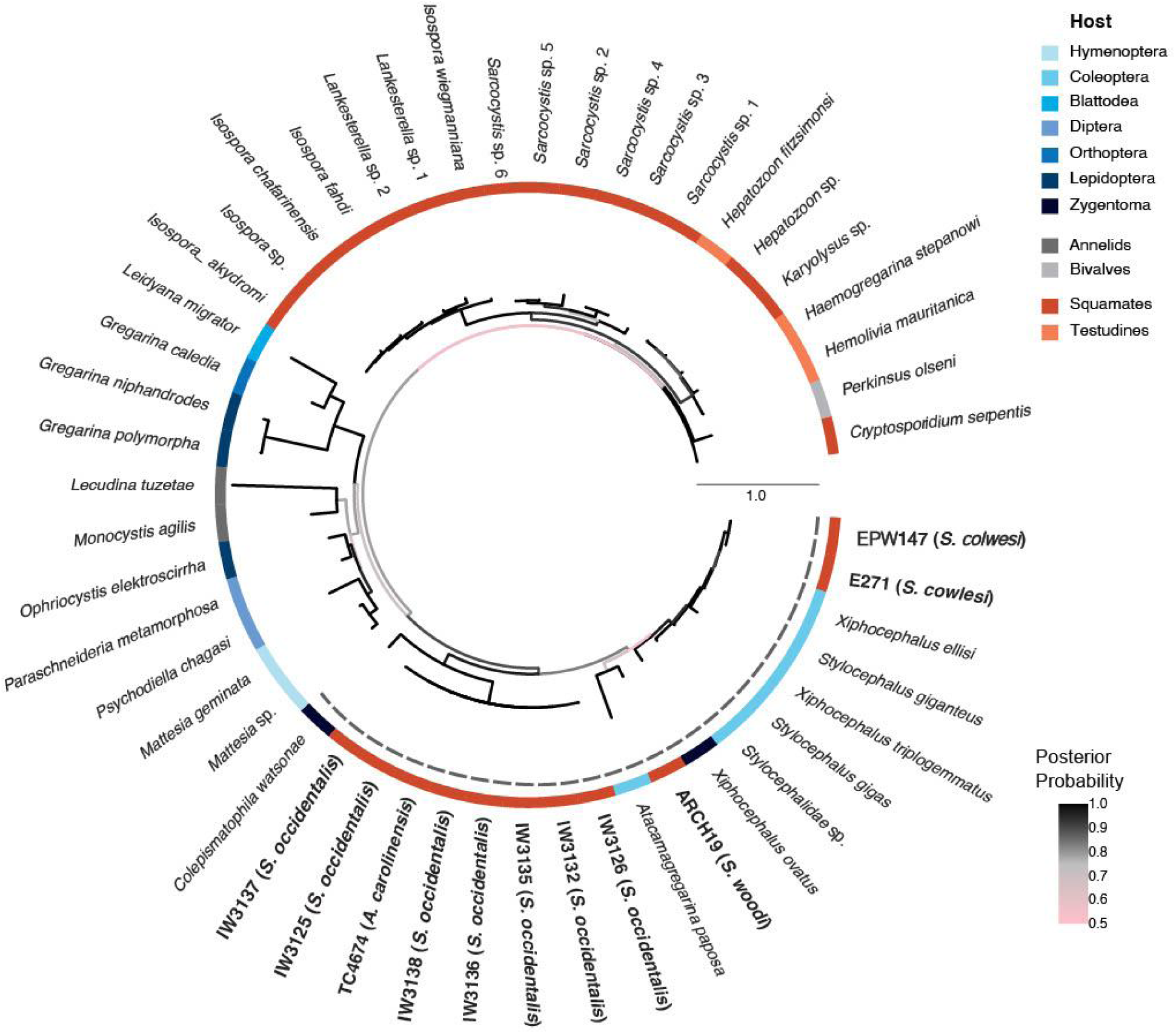
Pruned Bayesian phylogenetic tree of Gregarine-related sequences from this study and reference sequences. This tree is rooted with respect to pruned divergent clades. Sequences amplified from this study are in bold, and all accession numbers are in Supplemental Table 3. Posterior probability values were represented as a color gradient. Host taxonomic group is shown in the outer ring, colored by insect orders (blues), non-insect invertebrates (greys), and vertebrates (reds). The gray dotted line denotes the Stylocehalidae family. Branches that were distantly related to samples were pruned for improved readability (Supplemental Fig 1).

For our novel Perkinsea sequences, the sequence from sample MLY543 (*Anolis schwartzi* from Sint Eustatius) fell in a polytomy with environmental and frog-associated reference sequences from Africa and Europe (Fig. 3, Fig. S2). The sequences from samples IH473 (*Aspidoscelis exsanguis* from New Mexico) and R0013804 (*Anolis carolinensis* from Alabama) were sister to each other with a support value of 0.99 and sharing a most recent common ancestor with a Perkinsea freshwater environmental sample from Trenchford reservoir in the UK (Fig. 3) with a support value of 0.93. Both clades containing our sequences formed a polytomy with the parasitic genera *Parvilucifera*, *Cryoxcellia*, and NAG01 (Novel Alveolate Group 01). These lineages infect a diverse range of hosts, including marine dinoflagellates (*Parvilucifera*), Arctic fish (*Cryoxcellia*), and freshwater amphibians (NAG01), demonstrating the broad host diversity within Perkinsea. Because our sequences cluster among these known parasitic lineages, they may represent previously unrecognized parasitic associations in reptiles.

**Figure 3:**
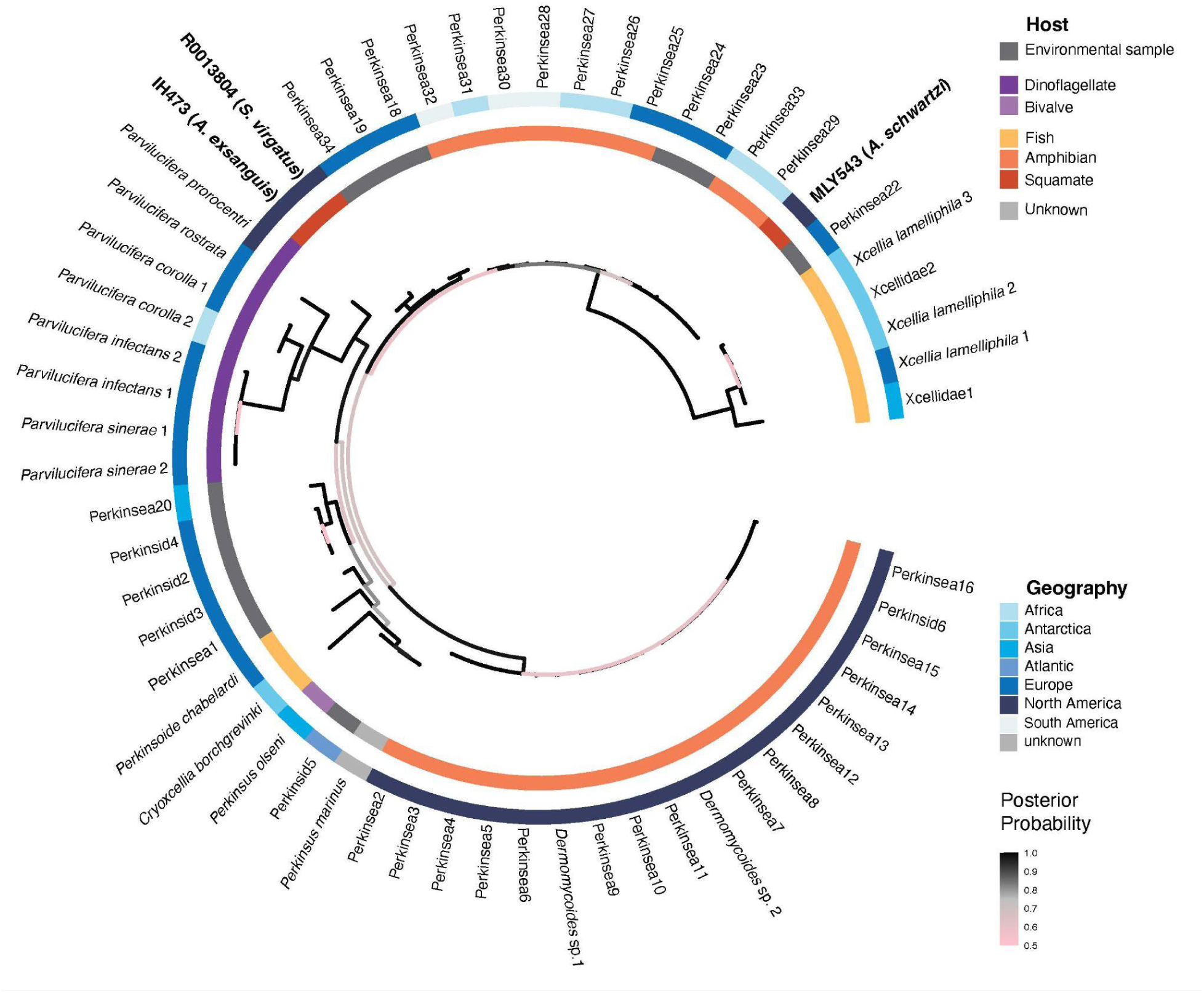
Pruned Bayesian phylogenetic tree of Perkinsea-related sequences from this study and reference sequences. This tree is rooted with respect to pruned divergent clades. Sequences amplified from this study are in bold with host species in parentheses, and all accession numbers are in Supplemental Table 4. Posterior probability values were represented as a color gradient. Host taxonomic group is shown in the inner ring and geographic origin is shown in the outer ring. Branches that were distantly related to samples were pruned for improved readability (Supplemental Fig 2).

## DISCUSSION

Our phylogenetic analyses placed the novel sequences in the taxonomic groups Stylocephalidae and Perkinsea and discuss their evolution as microbes that could become emerging reptile pathogens under the right conditions. Stylocephalidae is a family of gregarine alveolates. Gregarines are a diverse group of obligate parasites that primarily infect the digestive tracts of various terrestrial arthropods via ingestion of oocysts, the thick-walled zygote life stage (Clopton, 1999; Clopton, 2009), by the host. Gregarine alveolates, including *Xiphocephalus*, a genus within Stylocephalidae, have been found in soil, providing evidence that they could be spread as oocysts when detritivorous insects consume feces, corpses, or leaves (Yu et al., 2024). Stylocephalidae are known to infect many common arthropod species, including beetles, cockroaches, and silverfish, but their taxonomy and diversity are not well established (Clopton, 2009). For example, only three species from the genus *Stylocephalus* have DNA sequence data available (Mondal & Modak, 2025), even though every species of arthropod could potentially have a coevolved gregarine (Rueckert et al., 2019). This lack of molecular data poses a significant barrier to understanding phylogenetic relationships within this family and to detecting potentially pathogenic members of this clade in hosts. Additionally, to our knowledge, vertebrates have not been screened for the presence of these organisms, limiting our understanding of whether these organisms are solely associated with arthropods or found more broadly.

Perkinsea is a phylum of parasitic alveolates, comprising four known classes found in marine and freshwater environments. Perkinsea infects a range of organisms, including dinoflagellates, mollusks, fish, and amphibians (Itoïz et al., 2022). Marine Perkinsea taxa have been found infecting mollusks since 1946 (Itoïz et al., 2022), but freshwater Perkinsea appear relatively newly emerged (Chambouvet et al., 2014). The first report of freshwater Perkinsea infection was from a southern leopard frog (*Rana sphenocephala*) mortality event in 2003 (Davis et al., 2007). This parasite lineage can cause Severe Perkinsea Infection (SPI), causing abdominal distension, subcutaneous edema, skin discoloration, tissue destruction (Isidoro-Ayza et al., 2019), and yellowing and enlargement of the liver in tadpoles (Chambouvet et al., 2015) and, more rarely, in adult frogs. As Perkinsea has spread and caused mass amphibian mortalities in the US and UK, Perkinsea has become recognized as a widespread and highly deadly amphibian pathogen (Chambouvet et al., 2015). Infection occurs when free living zoospores are ingested or enter through mucous membranes, followed by penetration of the host’s tissues, where they develop into trophozoites. The trophozoites proliferate throughout the host’s tissues, leading to organ failure and mortality. After the host dies, the trophozoites develop into hypnospores with thick walls and remain in the sediment. When the hypnospores reenter water, they sporulate into zoospores, and the lifecycle repeats. As far as we are aware, terrestrial vertebrates other than amphibians have not been screened for Perkinsea. However, since this is an emerging area of research, unpublished screenings of Perkinsea in terrestrial vertebrates may exist.

The biology and host associations of the parasitic alveolates are poorly understood outside a few taxa. Therefore, investigating the presence of alveolates in novel host species is important for understanding their potential to emerge as significant pathogens (Swapna & Parkinson, 2017), as demonstrated by the recent mass mortality events of tadpoles caused by Perkinsea (Isidoro-Ayza et al., 2017). Furthermore, understanding host-microbe relationships are essential for accurate community characterization in microbial ecology. Perkinsea has a diverse host range, including dinoflagellates, mollusks, fish, and amphibians. Host specificity and range in this group are an emerging area of research (Chambouvet et al., 2014), and our results suggest that this research area should explore potential interactions with reptiles. Similarly, our results identified a second potentially novel interaction with gregarines related to the family Stylocephalidae, which are previously unreported in reptiles. Stylocephalidae gregarines have primarily been documented as parasites of arthropods, particularly darkling beetles (Clopton, 2009), which are consumed by many insectivorous lizards. The presence of Stylocephalidae-related DNA in reptile tissues raises the possibility of parasite spillover and a potential novel emergence of parasites into reptiles.

The presence of Perkinsea and Stylocephalidae in squamate tissues has a few possible explanations. The first is that these occurrences may represent dead-end infections, in which microorganisms infect one host but fail to transmit between hosts (Maia, 2015). In this case, it is possible that the Perkinsea or Stylocephalidae entered via consumption of infected prey items, and the infection went systemic, but could not be transmitted to other individuals. This outcome would be notable because it would show that the parasites have the capacity to transition out of the digestive tract and become a systemic infection. Another explanation is that these are novel parasites of reptiles that can transmit between hosts and have not yet been recorded. The final option is environmental contamination of the tissues. Perkinsea is present across a wide geographical range of freshwater (Chambouvet et al., 2015), marine, and soil environments (Metz et al., 2023). Stylocephaliade have been found in soil (Yu et al., 2024), and oocysts and gametocysts are known to persist in soil environments (Clopton et al., 2016). Due to this, our findings could be false positives for marking these taxa as reptile parasites. However, this is contamination is unlikely since we screened 1095 reptile samples and these were the only positives we received in these alveolate clades.

In the case of Perkinsea in squamate tissues, we believe this constitutes a true infection, which would be significant, as Perkinsea has currently only been found in aquatic ecosystems. Given the wide range of organisms Perkinsea is known to infect (Itoïz et al., 2022), and since most of the species that tested positive for Perkinsea (*Anolis schwartzi, Aspidoscelis exsanguis, Sceloporus virgatus*) are found in xeric environments, and are not known to submerge themselves in water (AuMArc & Maillot 2012, Pajarito Environmental Education Center 2025, Thomas C. Brennan 2023, Yuan et al. 2023), it seems likely that this is a novel parasite of reptiles as opposed to environmental contamination of samples. However, future testing should exclude the integument and only amplify parasite DNA from blood or tissue to eliminate the possibility of environmental contamination.

Since Stylocephalidae DNA was found in 11 samples from four different locations within the US, and clustered within the same clade with relatively high posterior probability, we believe this is a true infection. Since Stylocephalidae is found in beetles and silverfish that are common prey for lizards, it is likely that these microbes are able to enter lizard tissues through an ingestion pathway. Based on these results, we cannot say if lizards are a true host or a dead end host. However, it is also possible that environmental contamination caused these to be false positives. Stylocephalidae have been found in soil (Yu et al., 2024), and oocysts and gametocysts are known to persist in soil environments (Clopton et al., 2016). Since most of the samples were from tail sections, it is possible that soil was wedged between scales and could have given false positives. However, it is very difficult to extract DNA from an environmental gametocyst (Steele, et al., 2012), which may make detecting environmental contamination unlikely.

This paper is the first to establish the presence of the alveolate families Perkinsidae and Stylocephilidae in reptile tissues. Alveolata is a broad taxonomic group containing numerous understudied organisms. Freshwater Perkinsea has only recently been identified, and only in amphibians where it caused mass mortality events (Itoïz et al., 2022). Stylocephalidae has a variety of minimally studied species whose taxonomy is not well defined. Due to the lack of knowledge in these groups, but given recent work identifying herpetological infection-associated declines (Allain & Duffus, 2019)., further research is necessary to identify potential reptilian parasites. This study provides the first molecular evidence of Perkinsea and Stylocephalidae DNA in squamate tissues. Although infection cannot yet be confirmed, these findings highlight the need for targeted environmental sampling, histopathology, and experimental infection studies to clarify host range and transmission potential. Understanding the range of microorganisms in squamates will help in early detection of potential pathogens and allow preventative measures prior to the next panzootic disease in reptiles. Furthermore, expanding knowledge of Alevolata can expand our understanding of both current and future diseases.

## Supplemental figures

**Supplemental Table 1:** Sample information.

| State Sample was Acquired | Sample Name | Genus | Species | latitude | longitude | Collector Name | Year Collected |
| --- | --- | --- | --- | --- | --- | --- | --- |
| New Mexico | E271 | *Sceloporus* | *cowlesi* | 33.68476502 | -105.9200075 | Erin Westeen | 2021 |
| New Mexico | EPW147 | *Sceloporus* | *cowlesi* | 33.68476502 | -105.9200075 | Maggie Grundler | 2021 |
| Florida | ARCH 19 | *Sceloporus* | *woodi* | 27.18857 | -81.34875 | Iris Holmes | 2022 |
| California | IW 3125 | *Sceloporus* | *occidentalis* | 37.71865 | -119.6484 | Ian Wang | 2020 |
| California | IW 3126 | *Sceloporus* | *occidentalis* | 37.74297 | -119.43032 | Ian Wang | 2020 |
| California | IW 3132 | *Sceloporus* | *occidentalis* | 37.74303 | -119.43021 | Ian Wang | 2020 |
| California | IW 3135 | *Sceloporus* | *occidentalis* | 37.71851 | -119.37386 | Ian Wang | 2020 |
| California | IW 3136 | *Sceloporus* | *occidentalis* | 37.7183 | -119.37298 | Ian Wang | 2020 |
| California | IW 3137 | *Sceloporus* | *occidentalis* | 37.71843 | -119.37385 | Ian Wang | 2020 |
| California | IW 3138 | *Sceloporus* | *occidentalis* | 37.71975 | -119.64838 | Ian Wang | 2020 |
| Alabama | TC4674 | *Anolis* | *carolinensis* | 30.27888 | -87.68666 | Greg Pauly | 2012 |
| New Mexico | IH473 | *Aspidoscelis* | *exsanguis* | 32.35225 | 106.5573 | Iris Holmes | 2021 |
| Arizona | R0013804 | *Sceloporus* | *virgatus* | 31.91227 | -109.18478 | CUMV | 1999 |
| Sint Eustatius | MLY543 | *Anolis* | *schwartzi* | 17.50808 | -62.98565 | Michael Yuan | 2019 |

