## Supplemental Figures 1 & 2 for "Molecular evidence of the eukaryotic parasites Perkinsea and Stylocephalidae in lizard muscle tissue"

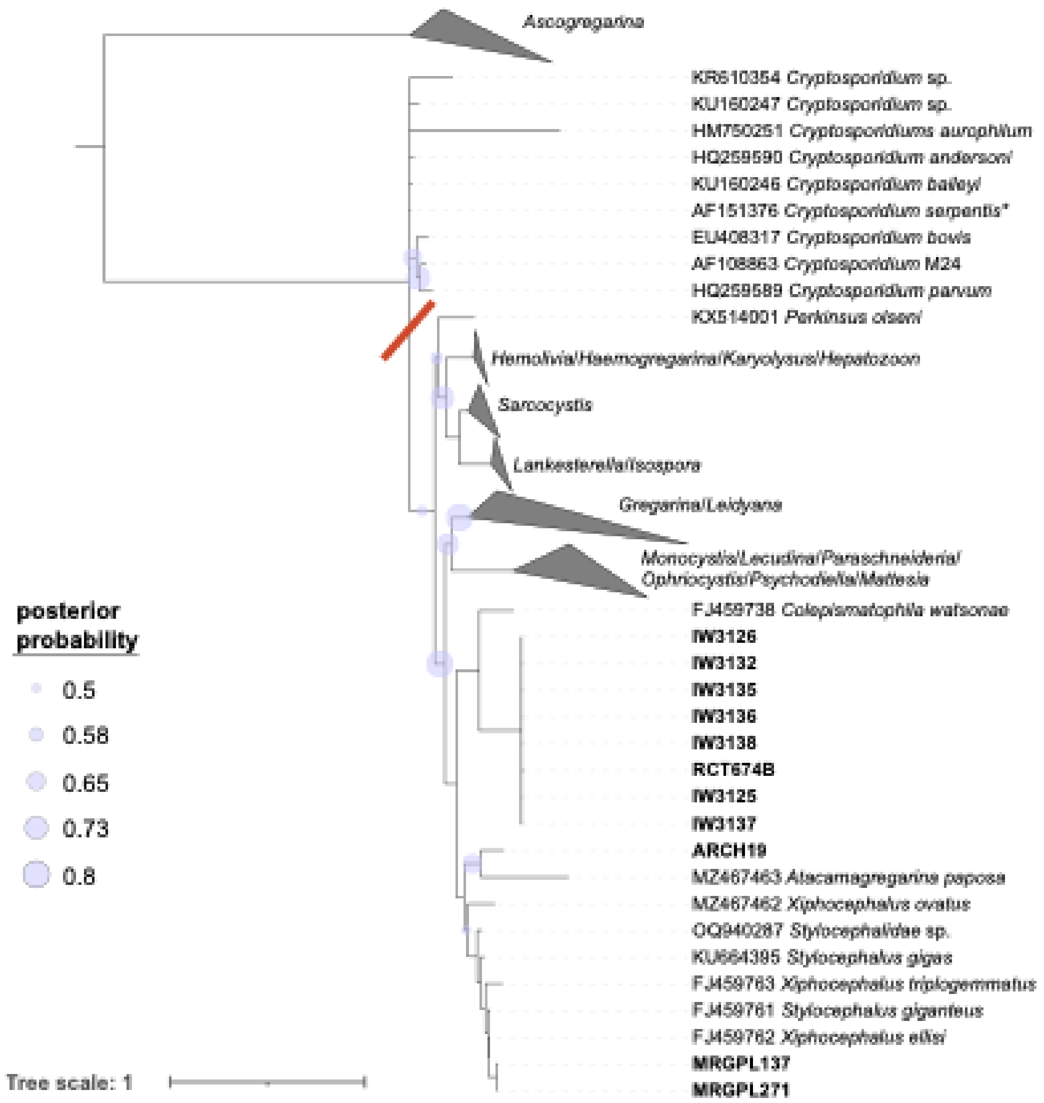


**Supplemental Figure 1.** Full (unpruned) gregarine phylogenetic tree with midpoint rooting. Taxa branching basal to the red slash, with the exception of one *Cryptospiridium* sequence (*), were pruned from the tree in Figure 2 for ease of visualization. Sequences from this study are in bold. Clades that are not closely related to sequences from this study were collapsed and are labeled by the genera they contain. Posterior probability values of 0.8 or less are shown at nodes, all other nodes have probabilities >0.8.

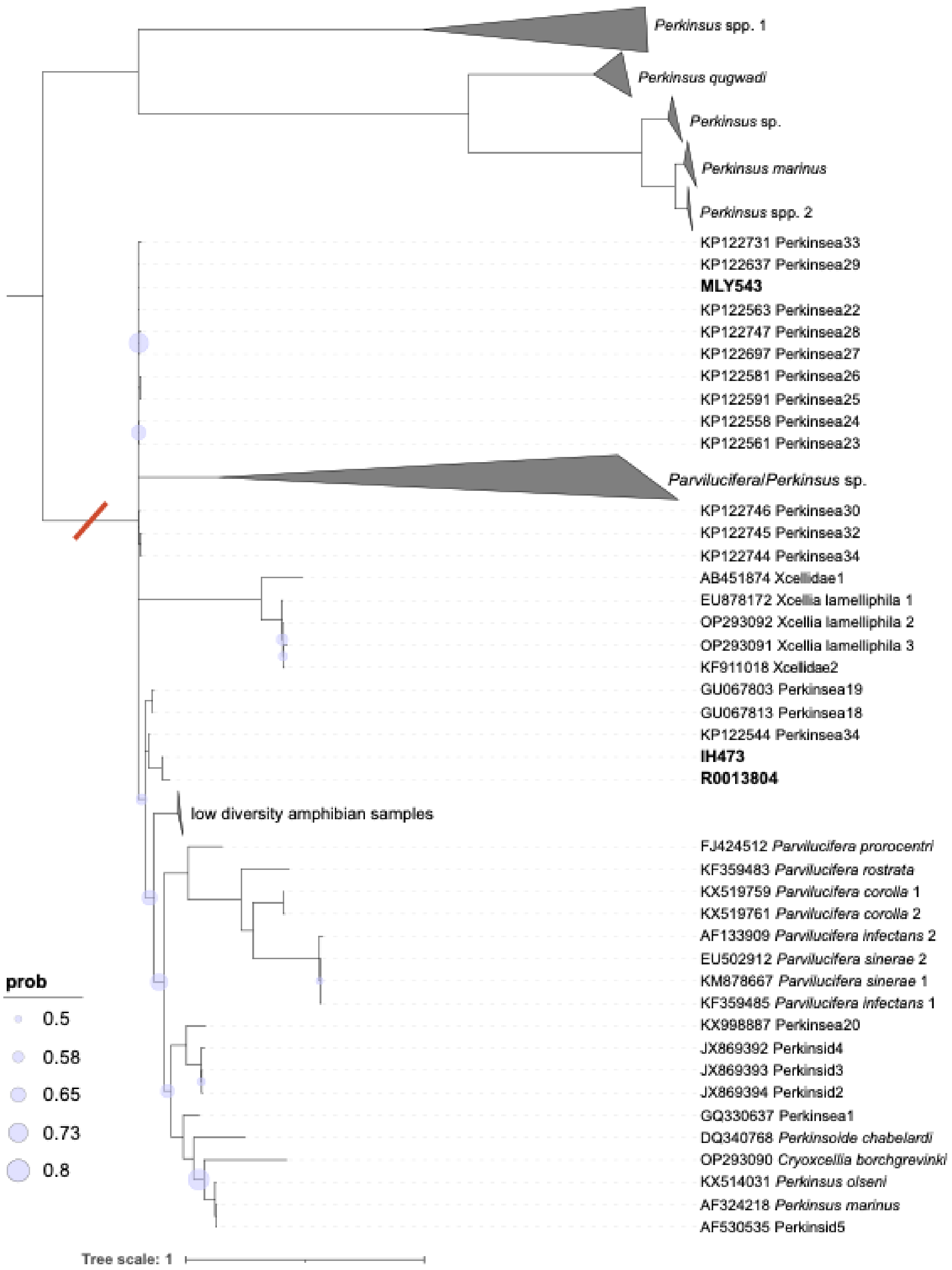


**Supplemental Figure 2.** Full (unpruned) *Perkinsea* phylogenetic tree with midpoint rooting. Taxa branching basal to the red slash were pruned from the tree in Figure 3 for ease of visualization. Sequences from this study are in bold. Pruned clades or large clades that are not closely related to sequences from this study were collapsed and are labeled as in Table S2. Posterior probability values of 0.8 or less are shown at nodes, all other nodes have probabilities >0.8.
